# Prevalence of water and nutrient availability in structuring fungal communities of healthy and decaying beech trees in forests

**DOI:** 10.1101/2025.05.27.656299

**Authors:** F. Fracchia, B. Dauphin, L. Walthert, R. Graf, R. Köchli, S. Pfister, A. Baltensweiler, A. Kohler, M. Peter

**Author notes:** These authors contributed equally (A. Kohler, M. Peter).

## Abstract

**Background and aims:** In forest ecosystems, trees interact with a broad range of soil microorganisms, such as ectomycorrhizal fungi, improving nutrition and water uptake and mitigating biotic and abiotic stress. In the context of the predicted more frequent and more severe droughts, it is critical to characterise how trees and their associated fungal partners respond to water shortage to improve future forest management.

**Methods:** We investigated the importance of diverse environmental predictors (e.g., edaphic, climatic, topographic) on the fungal communities associated with the root systems of decaying and healthy beech trees in natural beech forests. In parallel, we identified specific fungal taxa linked with their host vitality and water stress gradient.

**Results:** We observed that soil water shortage had a greater effect on the structuring of fungal communities than beech vitality or other environmental factors. We identified a core group of fungi that remained unaffected by water availability among all study sites, while other fungal species were more abundant in sampling sites more prone to water shortage. Finally, we showed that the abundance of ectomycorrhizal fungi was significantly associated with healthy beech root systems, while saprotrophic fungi prevailed in the roots of trees exhibiting severe decay.

**Conclusions:** Overall, our study highlights the major role of drought in structuring fungal communities associated with beech root systems and pinpoints the key fungal taxonomic groups found in healthy beech trees.

## INTRODUCTION

Forest trees are foundation and long-lived species that play an essential role in the structure and functions of the ecosystem. However, ongoing climate change alters the distribution of trees in two ways: first through progressive change in average climatic conditions (e.g. annual mean temperature, precipitation), and second through extreme climatic events. Biotic factors also contribute significantly to the tree mortality induced by drought, and the frequency and severity of pathogens attacks will rise with climate change (Hartmann et al. 2018; Patejuk et al. 2022). As the intensity and frequency of droughts and heat waves are expected to rise in many parts of the world (Dai, 2013; Hari et al. 2020; Pokhrel et al. 2021), assessing the fate of European forest tree species, beyond the biologically detrimental consequences, becomes a major socio-economic challenge (McDowell et al., 2020). In Europe, beech forests constitute economically and ecologically important ecosystems that are considered vulnerable to current and future climate changes, particularly in the French, German and Swiss lowlands (Gárate-Escamilla et al., 2019; Schuldt et al. 2020). Thus, the vulnerability of mature European beech to drought was evaluated based on a network of natural forest sites in Switzerland (Walthert et al., 2021), complementing previous findings on the drought resilience of beech saplings investigated under experimental conditions (Hagedorn et al., 2016; Pflug et al., 2018). Together, these studies raise questions about how this foundation species and its associated soil microorganisms, including their biotic interaction, will cope with extreme climatic events and other environmental changes.

Microorganisms associated with trees play a crucial role in the overall fitness and ecophysiological response of their host. Ranging from detrimental to beneficial, microorganisms participate in the biogeochemical cycling of nutrients, providing water and nutrients to their host, and offering protection against biotic (e.g. pathogens) and abiotic stresses (e.g. water stress, pollutants; Trivedi et al., 2020). At the soil–plant interface, it has been documented that the assembly of root fungal communities primarily depends on soil properties, providing the main reservoir for future root microbial communities (Beckers et al., 2017; Bonito et al., 2014; Mangeot-Peter et al., 2020; Uroz et al., 2016). Towards the root system, the rhizosphere is the first compartment where the host genotype starts to influence the microbiota composition through rhizodeposits (Reinhold-Hurek et al., 2015; Sasse et al., 2018). Finally, within the root endosphere, plant-microbe and microbe-microbe interactions are the main factors driving the assembly of the microbiota (Lareen et al., 2016; Trivedi et al., 2020; Veach et al., 2020).

The diversity and composition of soil microbial communities are also altered directly or indirectly by climate change. Drought have been reported to have both negative and positive effects on microbial species diversity, with different responses between bacterial and fungal communities (Hartmann et al., 2017; Preece et al., 2019). In addition, the impacts of drought conditions on plants indirectly affect microbial communities by altering root exudation (Gargallo-Garriga et al., 2018; Preece et al., 2021; Williams and De Vries, 2020) and litter inputs (Bardgett et al., 2008). These alterations in microbial species diversity and composition due to climate change may lead to a reduction in microbial activity, such as nutrient mineralisation and enzymatic activities, ultimately resulting in compromised functions and resilience of forest trees when facing a disturbed environment (Acosta-Martínez et al., 2014; Nguyen et al., 2018). Nevertheless, it remains unclear how biotic and abiotic factors shape soil microbial communities when the surrounding soil environment is disturbed, e.g. during water stress events. In particular, the role of ectomycorrhizal (EcM) fungi is not only known to confer beneficial partnerships for nutrient uptake and defence against pathogens (Martin et al., 2016), but is also supposed to support essential functions to mitigate detrimental effects of soil water shortage (Bogeat-Triboulot et al., 2004; Gehring et al., 2017).

Previous experimental works using drought treatments reported that fungal community composition was altered towards specific drought-tolerant associations of soil fungal species across water stress gradients (Anthony et al., 2021; Meisner et al., 2018; Winterfeldt et al., 2024) and depending on the host vitality (Hopkins et al., 2018; Jaeger et al., 2024). However, none of these studies investigated how water stress alters fungal communities associated with root systems in healthy and decaying beech hosts. Here, we explore the combined effect of edaphic, climatic, and topographic environmental factors on fungal communities associated with the root systems of healthy and decaying beech trees that suffered from the extreme drought in Central Europe in Summer 2018 (Schuldt et al., 2020). Using six natural beech forests across Switzerland, environmentally characterised in detail with soil moisture time series over 10 years and spanning a gradient of water availability, we asked the following questions: (1) Which environmental factors are the main drivers shaping fungal communities? (2) Do water stress and host tree vitality influence the structure, diversity, composition and assembly of the fungal communities? (3) Are there specific fungal taxa associated with the host tree vitality and the drought conditions in the soil.

## MATERIALS AND METHODS

### Study sites, in situ measurements, and defining the water stress gradient

The responses of fungal communities associated with beech root systems were investigated at six natural forest sites, scattered across Switzerland (Central Europe, approx. 46–48°N and 6–10°E). The sites are part of the forest soil database at the Swiss Federal Institute for Forest Snow and Landscape Research (WSL) holding data from combined soil and vegetation inventories (Walthert and Meier, 2017) and long-term time series of soil water potential (Walthert et al. 2021), leading a total of 158 environmental variables. A complete description of the methods used to measure these variables is provided in the supplementary methods. In brief, the six sites are located in regions with different climates between 530 and 880 m a.s.l., where mean annual temperature (1991–2020) ranged between 8.6 and 9.6°C depending on the site, and mean annual precipitation sum between 731 and 1117 mm (Table S1). The sites were covered by old-grown mixed deciduous forests, predominantly beech and oak, with a leaf area index (leaf area per unit of ground) between 3.6 and 5.8. Three sites were almost completely covered by a layer of herbs and shrubs, whereas these layers were sparse on the other sites. The soils on all sites had calcareous parent material and were therefore weakly developed, belonging to the A-C soil group, with high pH values throughout (measured in CaCl_2_), ranging from 5.6 to 7.7. In contrast to the pH values, the topsoil carbon, nitrogen, and phosphorus contents were highly variable between sites. The soils also showed a high inter-site variability in the storage capacity of plant available water (37-176 mm) due to the large variation in soil depth (30-180 cm), stone content (18-88 %) and soil texture (13-75 % clay and 8-57 % sand; Table S1). Roots were present down to the parent rock or down to the exploration depth because the soils had no barriers to root growth such as high soil density or oxygen deficiency.

To estimate the water stress status of beech trees at each site, we first measured the soil water potential (Ψ_soil_). Ψ_soil_ was measured hourly using MPS-2 dielectric sensors connected to EM-50 data loggers (Decagon Devices, Pullman, WA, USA). These sensors determine Ψsoil indirectly by assessing water content in their porous ceramic discs, relying on dielectric permittivity as an indicator. The recorded water content is then converted to Ψsoil based on the moisture release curve of the ceramic discs. MPS-2 sensors were placed within the same soil profile used for soil and root analyses, with two to three sensors embedded in the front wall at depths of 20 cm and 80 cm, and, where possible, at 140–160 cm and 180–200 cm. To enhance accuracy, Ψsoil measurements were temperature-corrected to 22 °C following the approach described by Walthert and Schleppi (2018). From these measurements, we used the Ψ_soil_ threshold of -800 kPa defined by Walthert et al. (2021) that is indicative for a high stress level in the trees. We calculated the integral corresponding to the stress threshold, as the integral is a composite index for water stress duration, intensity, and frequency (Myers, 1988; Wullschleger and Hanson, 2006). Here, we used the trapezoidal integration calculus with the *trapz* function from the pracma package.

We calculated the drought stress integral for all six sites for all soil horizons, hereafter defined as integrated drought index (IDI), and for both deep and topsoil separately, called Deep or Top Soil IDI, respectively. We tested whether the differences between IDI across the six studied sites are significant with a linear mixed effect model (*lmer* function from the lme4 package), with the aim to confirm a water stress gradient across the six studied sites.

### Tree roots sampling and preparation

For the six studied sites, we sampled 10 pairs of a healthy and a decaying understory beech tree that showed substantial damage after the 2018 summer drought. We aimed at sampling healthy and decaying beech pairs being in close vicinity and at the same elevation in steep sites. However, most sites had only few healthy trees and often no suitable decaying counterpart. Therefore, pairs of a similar stem diameter were searched for, but the stem diameter among pairs ranged between 0.5 - 8.5 cm. By counting the number of annual tree rings, we established the age of each beech tree ranging from 5 to 82 years old, with the youngest trees mainly present in site Neunkirch, while the oldest dominated in sites Chamoson and Büren. We carefully dug out the whole tree including the root system down to 30cm depth in a radius of 30 cm around the trunk. From each tree, five fine-root pieces (≤ 2 mm diameter) about 10 cm long were randomly sampled, pooled and placed in a plastic bag between moistened household paper. The bags were transported to the laboratory in a cooling box and stored at 4°C until processing. Within three weeks, sampled fine-roots were carefully washed under running tap water and placed in a petri dish with water under the binocular. From each root piece, one subsample of fine roots ≤ 1mm containing vital fine-root tips was randomly selected, dried on a paper towel, pooled in a 2 ml tube per sample, and stored at -20°C.

### DNA extraction and metabarcoding sequencing

The root samples were lyophilized and ground before DNA extraction using the sbeadex technology with a customised extraction protocol for plant material (LGC Genomics, Berlin, Germany) on an automated KingFisher 96/Flex instrument (ThermoFisher Scientific, Basel, Switzerland). DNA was dissolved in 100 µl AMB buffer (LGC Genomics, Berlin, Germany) and stored at -20°C until metabarcoding.

For metabarcoding of the fungal communities, we amplified the ITS2 region of the nuclear ribosomal internal transcribed spacer (ITS) region using primers 5.8S-Fun and ITS4-Fun developed by Taylor et al. (2016), which efficiently exclude nonfungal DNA templates. Primer sequences contained adaptors as indicated by the sequencing company (Génome Québec Innovation Center, Montréal, Canada). We used the following conditions for PCRs: For a total volume of 25 µl, we used 3 µl of a 1:10 dilution of DNA, the GoTaq G2 Hot Start Taq Polymerase (Promega, Dübendorf, Switzerland) with 2.5 mM MgCl_2_, 0.8 µM primers, 0.4µM dNTPs, 0.6 µg/µl BSA and 1.25U G2 Polymerase. The cycling conditions were 2 min at 95°C of initial denaturation, 34-37 cycles of 40 sec at 94°C denaturation, 40 sec at 58°C annealing, and 1 min at 72°C elongation, and a final elongation of 10 min at 72°C. We optimised the number of cycles for equal band intensities, which were visually checked by electrophoresis on a 1.5% agarose gel in 1X TBE buffer as suggested by Castaño et al. (2020) and Clemmensen et al. (2016). As quality controls, we included both negative controls and known fungal communities (mock communities) in PCRs and sequencing. The sequencing company (Génome Québec Innovation Center, Montréal, Canada) performed the indexing, normalisation, library construction and sequencing on a PacBio Sequel II system with the non-purified PCR products.

### Bioinformatics and statistical analyses

All the data analyses, statistics and graphical representations in this study were computed in R v.4.3.0 (R Core Team. 2023), using RStudio v.2023.03.1 (RStudio Team, 2023). All the figures were created using ggplot2 v.3.4.4 (Wickham, 2016). The analysis of beta diversity of fungal communities was performed using the *vegan* package v.2.6.4 (Oksanen et al., 2022). The linear mixed effect models (LMM) and the generalised linear models (GLM) were performed using the *lmer*, and glm functions, respectively, from the lme4 package v.1.1.35.1, while generalised linear mixed effect models (GLMM) were performed using the *gamm* function from the mgcv package v.1.9.0. Except when mentioned differently, these models were generated using the formula: Y ∼ IDI * Health * Age + (1 | Pairs), where IDI, Health and Age were considered as fixed factors and Pairs as random factor. Finally, elastic net models were generated using the *glmnet* function from the glmnet package and cross-validated with the *cv.glmnet* function.

### Sequence processing

After quality controls, sequence demultiplexing and barcode removal, fungal sequences were processed at the Genetic Diversity Center of ETH Zurich. Fungal amplicons were clustered into zero-radius Operational Taxonomic Units (zOTUs) using the UNOISE3 algorithm with 99% identity, corresponding to the ITS species level (Lindahl et al., 2013). Fungal sequences not assigned to the ITS region, after using the ITSx filter, were removed. Fungal sequences were affiliated using SINTAX with the appropriate UNITE.v9 Fungal Database reference (Nilsson et al., 2019). zOTUs with a BLAST identity lower than 88% and BLAST coverage lower than 80% at the species level were considered as chimaeras and removed from the dataset. We rarefied fungal communities with a number of sequences to 2500 per sample. zOTUs not present in at least four replicates, were removed from the analyses.

### Identifying the environmental predictors that best explained the fungal community variance

To identify the best environmental predictors shaping fungal communities, we first scaled and centred all 158 environmental variables and used Principal Components Analysis (PCA) to characterise the main drivers shaping the distribution of the studied sites. For downstream analyses, we kept environmental variables that exhibited the highest contribution to the first three principal components (cos^2^ ≥ 0.70) or with major biological meaning related to water stress. To avoid multicollinearity among the predictors, we investigated the correlations between the retained environmental variables using Spearman’s correlation coefficients. Environmental variables that were highly correlated (*r* ≥|0.75|; cf. Dormann et al., 2013) with drought-related parameters were removed. Fungal communities were transformed with the Hellinger method and environmental parameters were scaled and centred to generate multivariate elastic net models, after assuring to retain a reasonable variation inflation factor below two (Zuur et al., 2009).

### Fungal community structure associated with beech roots

Using the fungal community transformed with the Hellinger method, the heterogeneity of the compositional data was calculated via a detrended correspondence analysis (DCA) to decide whether linear Redundancy Analyses (RDA) or unimodal Canonical Correspondence Analysis (CCA) ordination methods are more suitable (Borcard et al., 2011). As the longest DCA ordination axis was below three, we performed a transformation-based RDA (tb-RDA) analysis using the *rda* function from the vegan package. Differences in fungal community structure between sites, beech vitality, age, and their interactions were visualised through multiple regressions using tb-RDA (*rda* function in vegan package), implementing the four environmental parameters retained as explanatory variables and stratified with sites. The significance of the final model with the selected environmental predictors on the fungal community was assessed using a permutational multivariate analysis of variance (PERMANOVA) with 1,000 permutations (*adonis2* function in vegan package), stratified with sites.

We used the same constrained analysis (tb-RDA) to investigate the relationships between fungal species and associated guilds with the environmental variables retained, the vitality and the age of the host. Variance Partition Analyses (VPA) was performed to quantify the proportion of explained variance for each environmental, age and health predictors using variancePartition package version 1.32.3 (Hoffman and Schadt, 2016).

### Fungal diversity and guilds associated with beech roots

We characterised the alpha diversity through measuring four indices: species richness, Shannon and Simpson diversity indices and Pielou’s evenness. The two databases FUNGuild (Nguyen et al., 2016) and FungalTraits (Põlme et al., 2020), were combined to classify each fungal taxon, from the order to the species levels, into an ecological trophic guild. A confidence threshold was applied to only keep “highly probable” and “probable” affiliated trophic guilds, whereas remaining zOTUs were assigned as “unknown”. We characterised the effect of water stress, beech vitality, age and their interactions on fungal richness and diversity using LMM and GLMM (beta distribution), with beech pairs as random effect. Finally, we assessed the influence of water stress, host vitality, age and their interactions on the relative abundance of fungal guilds using GLMM with a beta distribution, again with beech pairs set as random effect.

### Fungal species associated with a water stress gradient and beech vitality

To identify specific fungal taxa significantly associated across a water stress gradient and between host vitality, linear discriminant analysis (LDA) effect size (LEfSe), (Segata et al., 2011) was used through the microeco package v.1.4.0. A threshold of LDA value below or equal to two was used. These results were cross-validated with another approach based on Indicator Species value (Cáceres and Legendre, 2009; Dufrêne and Legendre, 1997) using the indicspecies package v.1.7.14. We only considered fungal species that showed significant associations in both analyses.

### Stochasticity analysis

Finally, the stochasticity resulting in the assembly of fungal communities associated with healthy and decaying beech root systems across the water stress gradient was quantified with the Normalised Stochasticity Ratio (NST), using the NST package v.3.1.10 (Ning et al., 2019). Fungal communities were transformed using the Hellinger method and both taxonomic (Jaccard, Bray-Curtis) and phylogenetic (UniFrac, weighted UniFrac) distances were used. Variation of NST was bootstrapped 1000 times and relationships between NST and water stress gradient depending on beech vitality were assessed using GLMM with a beta distribution, and an Anova test was performed to evaluate the significance of host health and water stress on fungal communities’ assembly processes.

## RESULTS

### Defining a water stress gradient among the studied sites using soil water potential (Ψ_soil_)

To investigate the influence of water stress on the fungal communities associated with beech root systems, we characterised the intensity of water stress for the beech trees among the six studied sites. We observed significant differences of IDI between the studied sites, from Tamins as the “wettest” site and Chamoson as the “driest” site for the downstream analyses (LME, P < 0.05; Figure S1). The IDI greatly varied over the years and seasons, with 2018 being the driest year. The IDI ranged from 1043.3 to 1814.4 area under the curve (AUC) from the driest site Tamins to the wettest site Chamoson. Finally, the maximal IDI over 20 consecutive days ranged between 28.26 AUC for the driest site to 14.44 AUC in the wettest site. Overall, these findings substantiate the presence of a water stress gradient among the six studied sites.

### Prevalence of key environmental factors rather than host age and health contributing to the structure of fungal communities

After carrying out multivariate elastic net models, we retained four environmental factors explaining most of the variance between fungal communities (i.e. the IDI, the Deep Soil IDI, Richness of the Arboreal Layer (RAL), and organic phosphorus (P.org)).

We used these four environmental predictors, in addition to the age and vitality of the beech hosts, to constraint statistical models of the fungal communities associated with the root systems of healthy and decaying beech trees across a water stress gradient. Through tb-RDA, we observed a clear separation of fungal communities depending on study sites rather than on the citality of their host (Figure 1, Table 1). The first two axes explained 10.79 % of the total variation in fungal composition (6.57 % and 4.22 % of the total variance for tb-RDA1 and tb-RDA2, respectively). In addition, using linear regression and permutation tests, we identified fungal taxa significantly associated with each site (Figure 2). For example, the saprotrophic fungi *Pezicula radicicola* and *Mycena leptocephala* were significantly associated with the “driest” site Chamoson, while the endophyte *Ilyonectria mors-panacis* and the EcM *Sebacina cystidiata* were correlated with the “wettest” sites Neunkirch.SH.Ei and Tamins (Figure 2).

**Figure 1.**
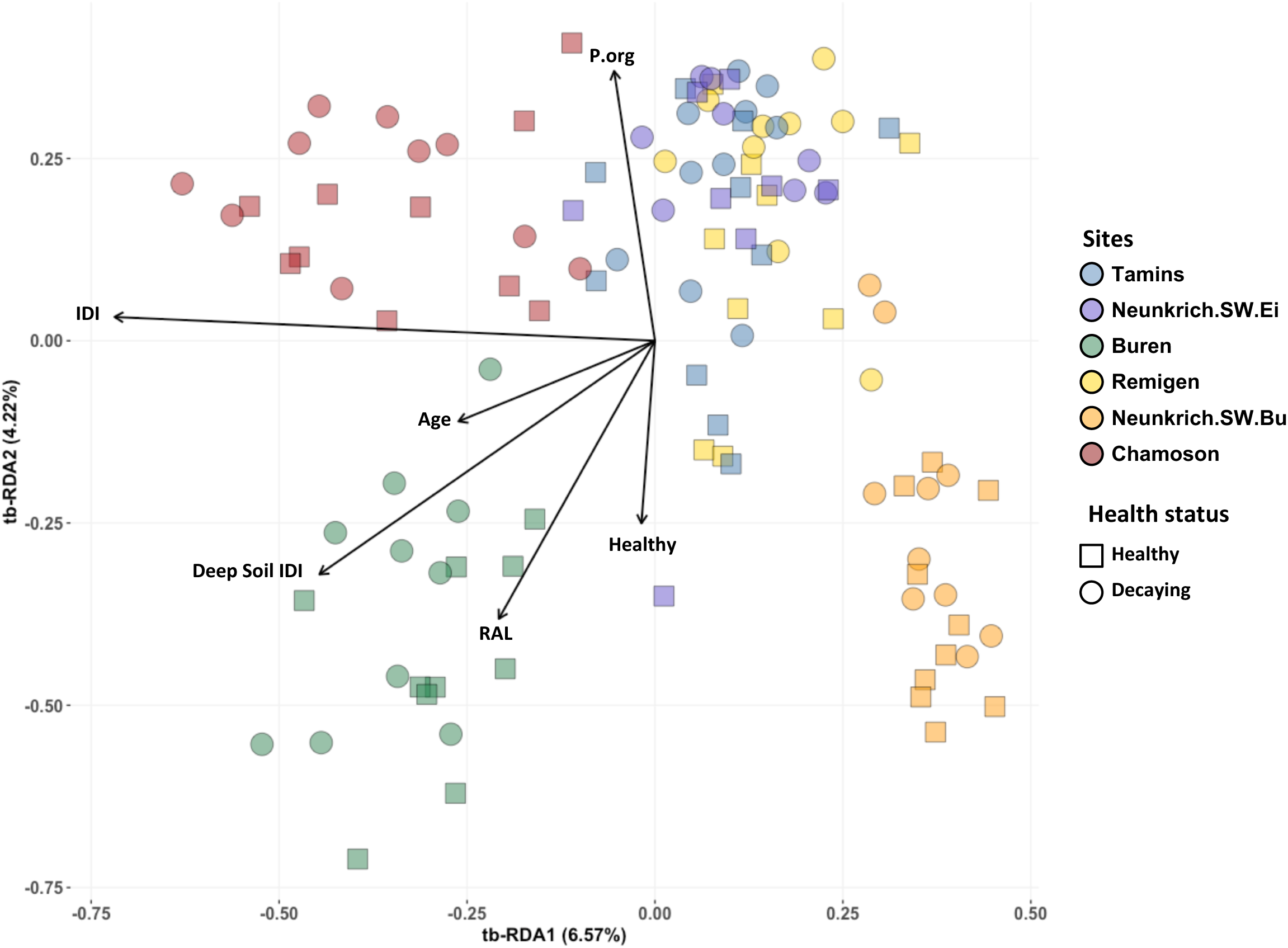
Redundancy analysis (RDA) showing distribution of the fungal communities according to environmental predictors, age and beech vitality. Vectors indicate significant variables constraining the microbial communities after multiple regression analyses and 1000 permutations (n = 8-10, FDR corrected, p.adj ≤ 0.01). The significant effects of environmental variables and beech vitality were assessed by PERMANOVA after 1000 permutations (p.adj ≤ 0.01). Colors indicate the sites and shape relate to the vitality of the beech host.

**Figure 2.**
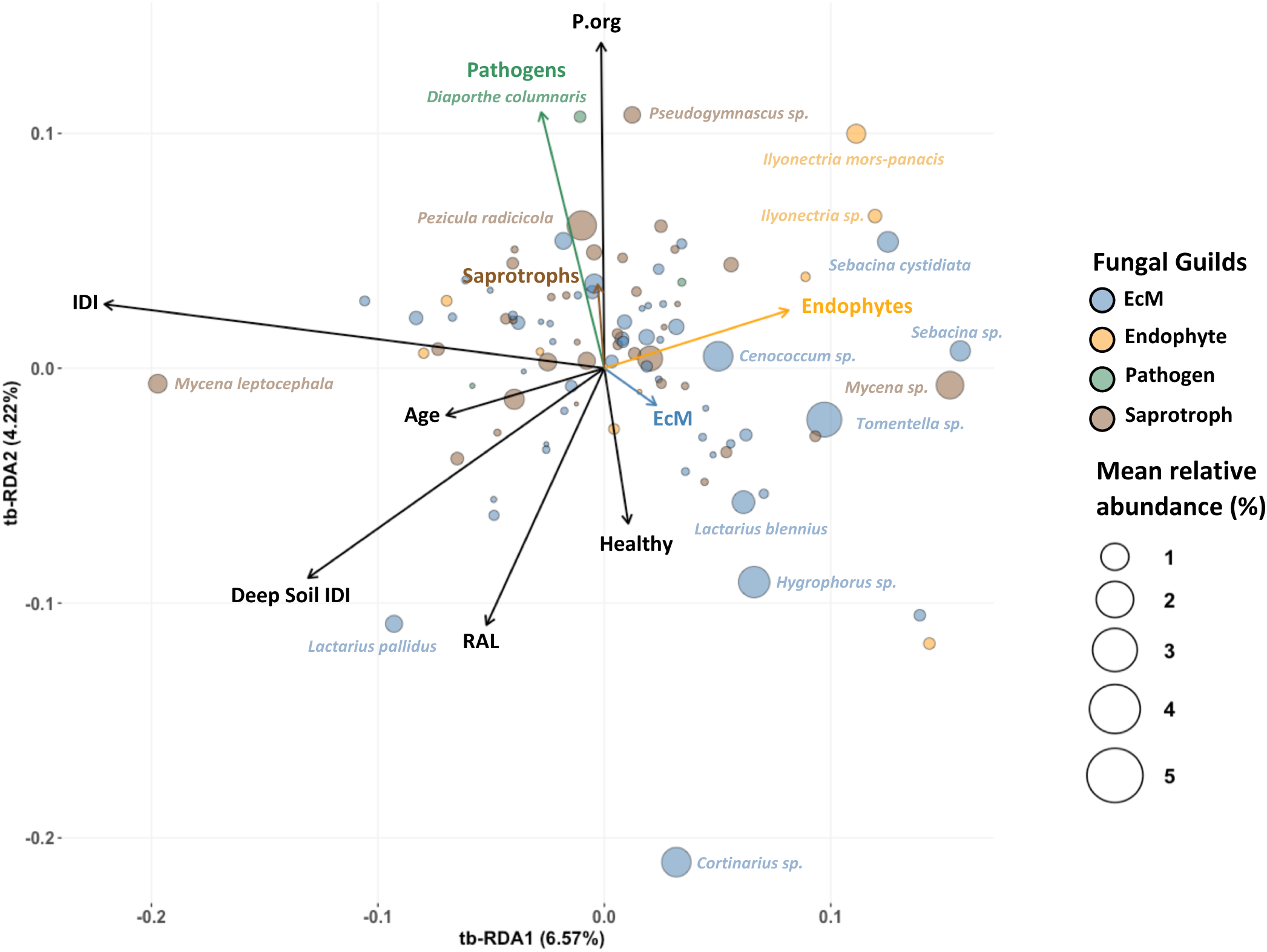
Redundancy analysis (RDA) showing distribution of the fungal species and guilds according to environmental predictors, age and beech vitality. Vectors indicate significant variables constraining the microbial communities after multiple regression analyses and 1000 permutations (n = 8-10, FDR corrected, p.adj ≤ 0.01). The significant effects of environmental variables and beech vitality were assessed by PERMANOVA after 1000 permutations (p.adj ≤ 0.01). Colors indicate the fungal trophic guilds and the size of the circle the relative abundance of the fungal species.

The permutational multivariate analysis of variance (PERMANOVA) confirmed the significance of the model and all environmental variables investigated (adjusted P < 0.01). Overall, the environmental variables, the vitality of beech trees and their age explained 24.4% of the variation in fungal community composition across sites, while 75.6% remained unexplained (Table 1). Using PERMANOVA and variance partition analyses (VPA), we showed that the environmental conditions associated with each site contributed 19.8% of the variance of the fungal communities, which is significantly more than the age and the vitality of the beech host (3.5% and 1.2%, respectively; Figure S2, Table 1). Noteworthy, drought-related factors IDI and Deep soil IDI with 6.8% and 5.7% of the variance were the major ones structuring fungal communities, rather than soil properties (P.org: 3.8%) or vegetational such as richness of the arboreal layer (RAL: 3.5%) (Figure S2, Table 1).

Finally, we found that the EcM guild correlated with healthy hosts, while saprotrophs and pathogens were associated with decaying beech and P.org (Figure 2). These findings were further supported by GLMM analyses (see below).

### Influence of environmental factors, age and host vitality on fungal richness and diversity

We first assessed the alpha diversity of these communities to characterise the influence of the water stress gradient on fungal communities associated with the root systems of healthy and decaying juvenile beech trees. A total of 1,361 fungal OTUs were identified from 669,142 high-quality sequences at 99% similarity level, leading to 311 fungal species after applying different quality filters. We did not observe any significant differences in richness or diversity (Shannon entropy, Simpson index and Pielou’s evenness), neither at the fungal OTUs nor at the species taxonomy level across the water stress gradient and beech age (n = 8-10, LME, P > 0.05; Figure 3, Table S2). Conversely, the vitality of the beech host had a significant effect on OTUs and fungal species richness but not diversity, with an increase between healthy- and decaying-associated communities (n = 8-10, LME, P < 0.05; Figure 3, Table S2).

**Figure 3.**
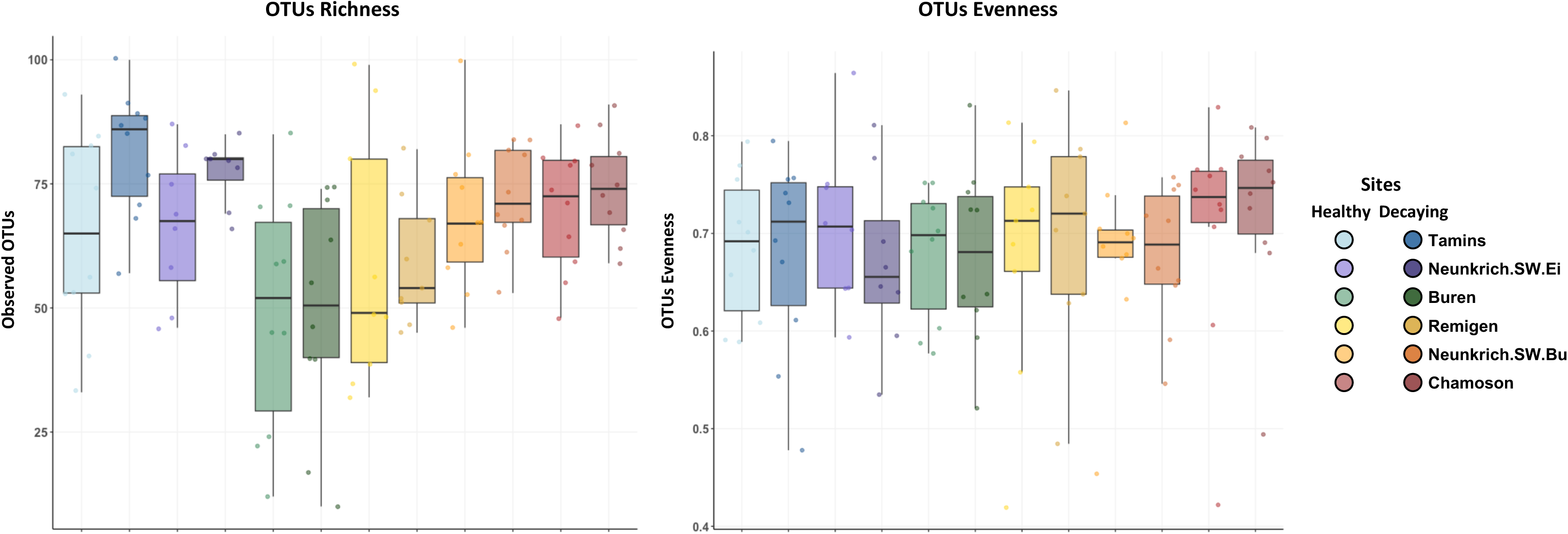
Fungal OTUs richness and diversity (Pielou’s evenness) between studied sites, the age and the vitality of the beech host. Fungal richness (left panel) and diversity (Pielou’s evenness; right panel) were not altered by water stress gradient, nor by the age of the beech host (n = 8-10, LMM and GLMM (beta distribution), p.value > 0.05). However, the richness of fungal OTUs significantly increased between healthy and decaying beech host (n = 8-10, LMM, p.value ≤ 0.05).

### Influence of environmental factors, age and host vitality on community composition

Looking at the overall composition of the one percent most dominant fungal species in these communities, we visually identified three groups showing similar abundances across the water stress gradient, without a clear delineation of the vitality of their host (Figure S3, Table S3): (1) fungal species that were present in all study sites (e.g. *Cenococcum* sp., *Penicillium* sp., *Cladiophialophora* sp.), (2) fungal species that tended to decrease (e.g. *Trichoderma* sp., *Metarhizium* sp., *Pseudeurotiaceae* sp.) (3) or increase (e.g. *Lactarius blennius*, *Oidiodendron chlamydosporicum*, *Absidia* sp.) with increasing water stress (Figure S3, Table S3).

Overall, the fungal communities on beech roots were dominated by EcM fungi (40.98 ± 0.26%), followed by saprotrophic fungi (27.41 ± 0.15). Only few endophytes (4.24 ± 0.08%), and pathogens (0.98 ± 0.05%) were detected, with a quarter of species whose trophic mode remains unknown (26.39 ± 0.14%) (Table S4). We observed a significant decrease of EcM relative abundance between the root systems of healthy and decaying juvenile beech (GLMM, P < 0.05; Figure 4, Table S4). On the contrary, the relative abundance of saprotrophs increased significantly from the root systems of healthy to decaying beech (GLMM, P < 0.05; Figure 4, Table S4). Concerning the influence of the water stress gradient, we found no effect on the relative abundance of any fungal guild (GLMM, P > 0.05).

**Figure 4.**
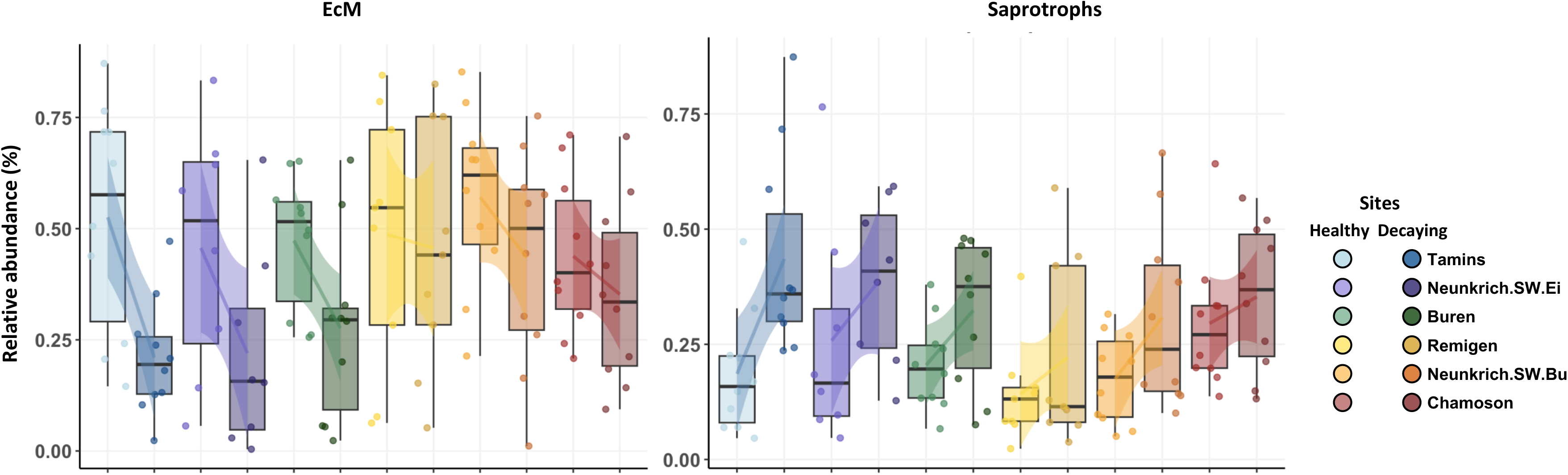
Relative abundance of fungal guilds depending on sites and beech vitality. The relative abundance of EcM significantly decreased while the relative abundance of saprotrophs significantly increased between the root systems of healthy and decaying beech (n = 8-10, GLMM (beta distribution), p.value ≤ 0.05), but were not altered by water stress nor the age of the beech host (n = 8-10, GLMM (beta distribution), p.value > 0.05).

At the species level, fungal communities were dominated by the EcM fungi *Tomentella* sp.*, Hygrophorus* sp., *Cenococcum* sp. and *Cortinarius* sp. (5.24 ± 0.89%, 4.12 ± 0.69%, 3.62 ± 0.47%, and 3.6 ± 0.93% relative abundance, respectively), as well as the saprotrophic fungi *Pezicula radicicola*, *Mycena* sp., and *Penicillium* sp. (3.54 ± 0.73%, 3.09 ± 0.72% and 2.46 ± 0.37%, respectively; Figure S3, Table S3).

### Specific fungal taxa are associated with water stress gradient and host vitality

Using linear discriminant analysis effect size (LEfSe), we identified 17 taxa significantly associated with the water stress gradient (Figure 5). Among them, the unidentified *Helotiales*, the saprotroph *Absidia*, and the endophytic genus *Oidiodendron* were significantly associated with the “driest” site Chamoson. Two EcM fungi, *Lactarius blennius* and *Sebacina* sp., were associated with the second driest site Neunkrich.SW.Bu (Figure 5). Interestingly, the abundance of these drought-associated fungal taxa was significantly positively correlated with the two water-stress related variables. On the contrary, *Mortierella amoeboidea*, the unidentified *Pseudogymnoascus* sp., and *Idriella* sp. were significantly associated with the two less drought-stressed sites Tamins and Neunkrich and negatively correlated with water stress related variables (Figure 5). Regarding the vitality of the beech host, three fungal saprotrophs were significantly associated with decaying hosts, namely *Pezicula radicicola*, *Hymenopellis radicata*, and *Camposporium appendiculatum* (Figure 6).

**Figure 5.**
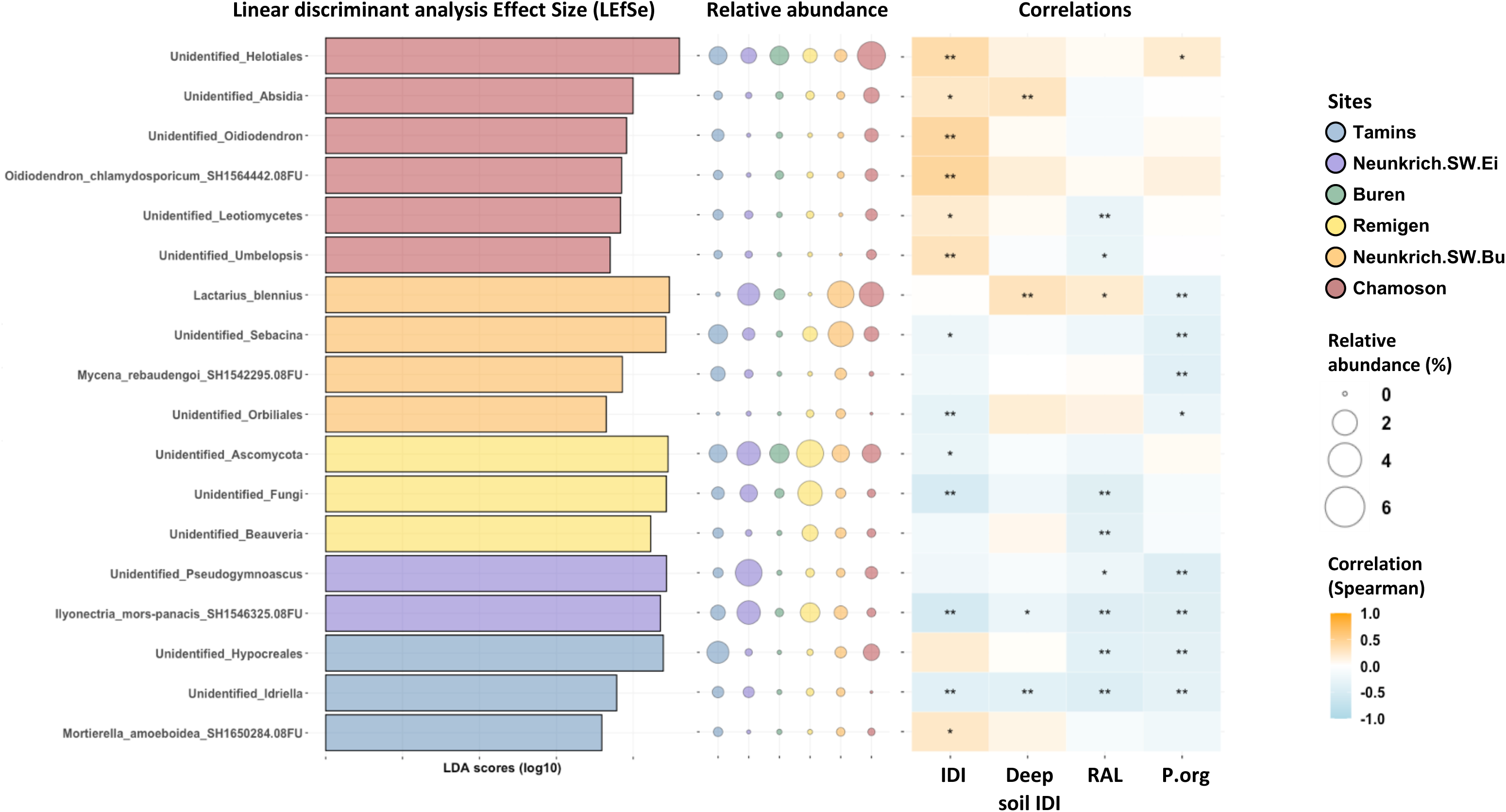
Linear discriminant analysis Effect Size (LEfSe) identified fungal species significantly associated with a water stress gradient. LDA scores, relative abundance of fungal species identified by LEfSe and their correlations with the 4 environmental predictors retained. Only species present at all sites were considered for the analysis. FDR corrections were applied to LDA scores.

**Figure 6.**
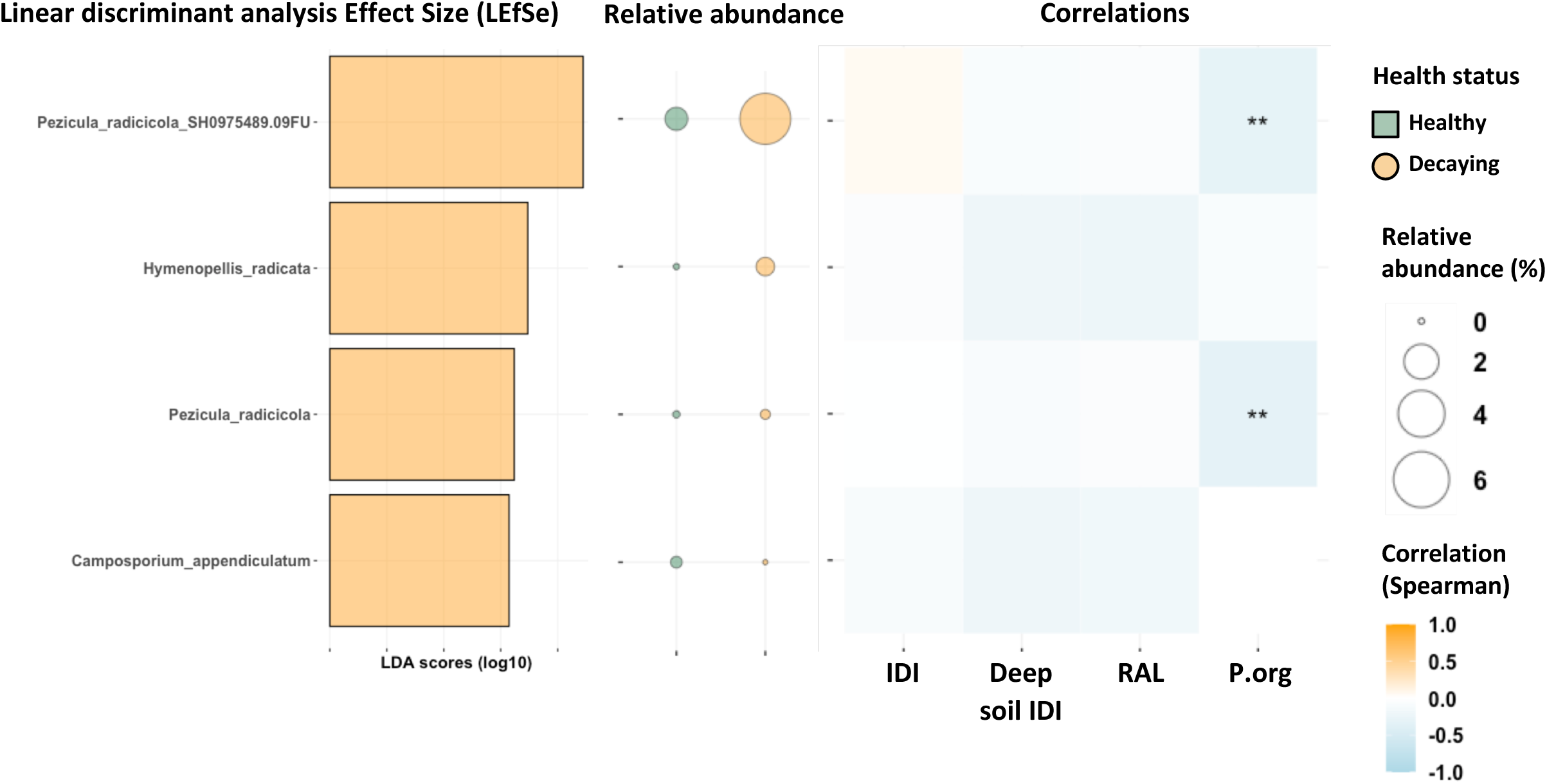
Linear discriminant analysis Effect Size (LEfSe) identified fungal species significantly associated with healthy and decaying beech root systems. LDA scores, relative abundance of fungal species identified by LEfSe and their correlations with the 4 environmental predictors retained. FDR corrections were applied to LDA scores.

### Importance of water stress and host vitality in the stochastic and deterministic assembly processes of fungal communities

Using the Normalised Stochasticity ratio (NST) developed by Ning et al., (2019), we found that stochasticity was the predominant assembly mechanism for the observed fungal communities (Figure 7). We observed a significant interaction between the water stress intensity and the host vitality, with deterministic processes being significantly more important in fungal communities associated with decaying beech hosts than healthy ones (GLMM, P < 0.05; Figure 7). Examining the assembly process depending on fungal functional guilds, we revealed that pathogenic and endophytic fungi exhibited a lower stochastic assembly than saprotrophs and EcM (Figure 7). Noteworthy, we observed a significant influence of the host vitality on the assembly processes of EcM, saprotrophs, and pathogens but not of endophytes (GLMM, P < 0.05; Figure 7): The stochasticity of EcM and saprotroph communities increased significantly with increasing water stress when associated with healthy beech while remaining stable in decaying beech. (GLMM, P < 0.05; Figure 7). On the contrary, for pathogenic fungi, the stochasticity of communities associated with healthy beech decreased significantly from the least stressed to the most stressed environment and remained stable in decaying beech (GLMM, P < 0.05; Figure 7). For endophytes, stochasticity increased over the water stress gradient for both communities on healthy and decaying hosts.

**Figure 7.**
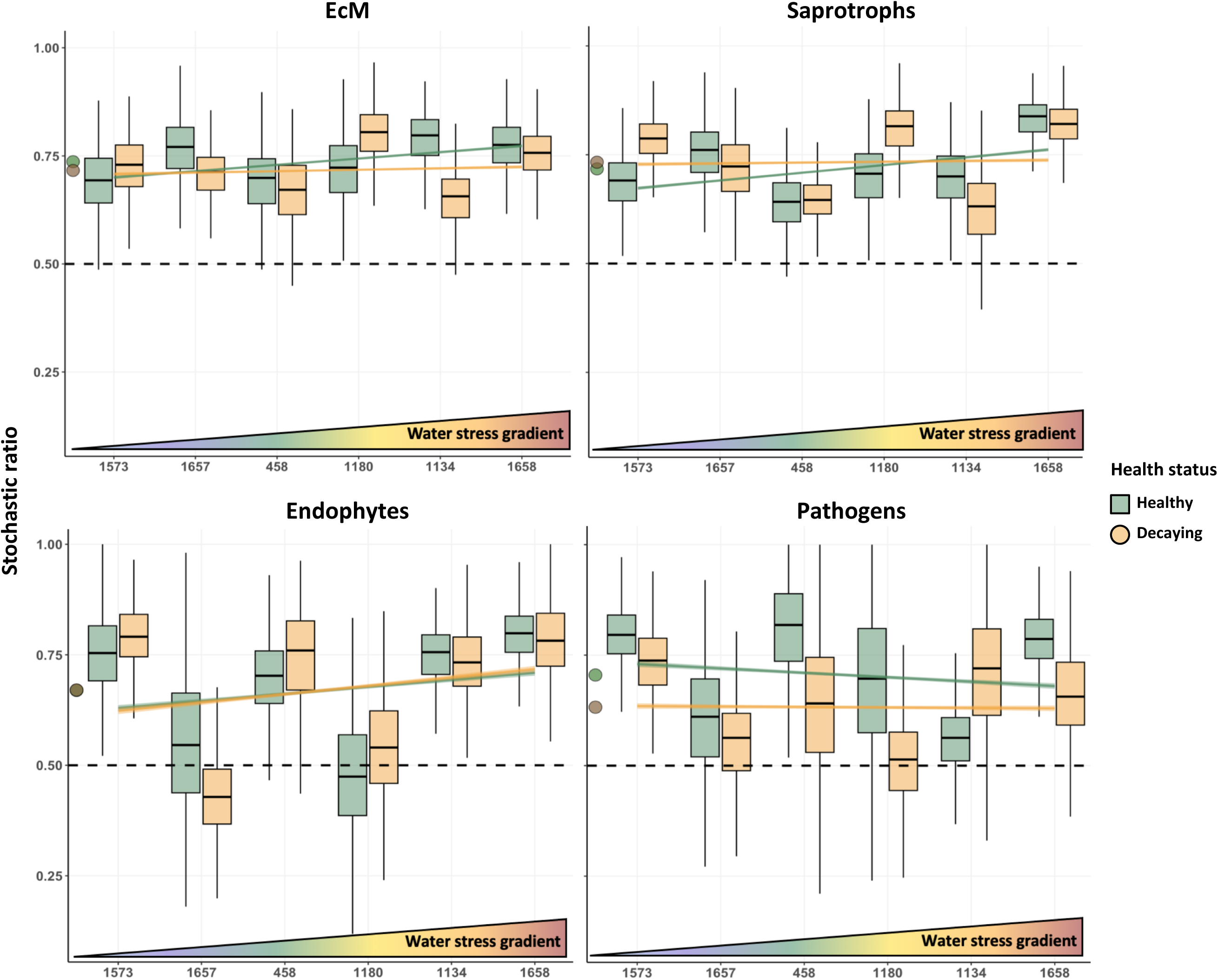
Normalised Stochastic Ration (NST) of EcM and saprotroph communities associated with the root systems of healthy and decaying beech across a water stress gradient. The stochasticity of the assembly process of EcM and saprotrophs communities increased along a water stress gradient in healthy host but remained unchanged in decaying beech. The vitality of the beech host and water stress had a significant effect on the assembly process of EcM and Saprotrophs (GLMM, P value ≤ 0.05). The stochasticity of endophytes increased significantly along a water stress gradient in both healthy and decaying hosts (GLMM, P value ≤ 0.05), and remained unaffected by the vitality of the host (p.value = 0.65). On the contrary, the stochasticity of the assembly process of pathogens decreased in healthy host along a water stress gradient but remained unchanged in decaying beech. The vitality of the beech host had a significant effect on the assembly process of pathogen communities (GLMM, P value ≤ 0.05). Black dashed line represent the cutoff between stochastic (> 50%) and deterministic (< 50%) assembly process. Circles indicate the overall mean and colors refer to the vitality of the beech host.

## DISCUSSION

It has been well documented that the assembly of root fungal communities is likely to depend on environmental factors, such as soil properties (Beckers et al., 2017; Bonito et al., 2014; Mangeot-Peter et al., 2020; Uroz et al., 2016), climatic conditions (Castro et al., 2010; Preece et al., 2020, 2019; Waldrop and Firestone, 2006), and the associated host trees (Cregger et al., 2018; Morella et al., 2020; Veach et al., 2020; Wagner et al., 2016). However, the main factors that shape fungal communities when all these factors are combined are still unclear. Indeed, very few studies have considered the mixed effect of environmental factors in shaping fungal communities during water stress (but see Anthony et al., 2021; Fu et al., 2022; Rillig et al., 2019; Zhou et al., 2020), and even fewer when integrating the vitality of the host in natural environments (but see Hopkins et al., 2018; Jaeger et al., 2024).

Overall, understanding the environmental drivers that determine the assembly of microbial communities is of great interest for managing the future European forests in the context of environmental change. In this study, we revealed that (1) water stress rather than individual climatic factors or soil properties, or age and vitality of the host is the main predictor of the assembly of fungal communities associated with beech root systems, (2) the vitality of the beech host, but neither water stress nor age, is altering the relative abundance of EcM and saprotrophs, (3) specific fungal taxa are associated with water stress conditions and host vitality, and (4) the assembly process of fungal communities is altered by both water stress and vitality of the host in a guild-dependent manner.

### Prevalence of water stress in shaping fungal communities associated with healthy and decaying juvenile beech trees

Fungal communities are altered by a broad range of biotic and abiotic parameters. Regarding abiotic variables, Zhou et al. (2020) investigated, through an exhaustive meta-analysis of 1,235 soil microbial communities, the influence of global change factors (e.g. elevated CO_2_, warming, and P addition) and their combination on microbial communities. They concluded that the variation of pH caused by global change factors was the main driver of soil microbial diversity and functionality (Zhou et al., 2020), as previously suggested in other studies (Lagueux et al., 2021; Rousk et al., 2010; Yang et al., 2021). Yet, it remains unclear which environmental factors are the main drivers of microbial assembly (Anthony et al., 2021; Birnbaum et al., 2019; Delgado-Baquerizo et al., 2016; Hendershot et al., 2017; Li et al., 2023; Moore et al., 2021; Zuo et al., 2023). For example, another global meta-analysis of 325 soil communities showed that the drivers of microbial diversity often differ among studies (Hendershot et al., 2017). In addition, while it is well known that the plant species, genotype and therefore physiology play a major role in recruiting its microbiota (Cregger et al., 2018; Morella et al., 2020; Veach et al., 2020; Wagner et al., 2016), none of the cited studies considered the influence of the host vitality in shaping microbial communities. As European forests are predicted to suffer more and more from extreme climatic events with increasing tree mortality, and the microbiota participate in the extended phenotype of the host tree (Whitham et al., 2003), it becomes necessary to investigate the combined effect of environmental factors and host vitality in shaping the host-associated microbial communities. To our knowledge, very few studies have attempted to disentangle these combined biotic and abiotic factors, and these works only focused on soil microbial communities and, thus, neglected the importance of host selection in shaping its microbiota (Hopkins et al., 2018; Jaeger et al., 2024).

Here, we revealed that drought related factors were the main drivers shaping fungal communities. Integrated drought index (IDI), the IDI in the deep soil, the Richness of the Arboreal Layer (RAL), and the organic phosphorus content (P.org) were the most significant drivers of fungal community structure. The two drought-related variables IDI and Deep Soil IDI accounted for more than half of the total explained variance of fungal communities (i.e. 12.5%), while the host age and its vitality only accounted for 3.5 and 1.2%, respectively. Soil pH has been frequently reported as the major soil factor influencing microbial community (Glassman et al., 2017; Lagueux et al., 2021; Tedersoo et al., 2020; Yang et al., 2021), also in beech forests across Europe (Coince et al., 2014). In the present study, soil pH was not retained as one of the most important predictors of fungal structure. This observation can be explained as the pH was more homogeneous, ranging from 5.6 to 7.2, than other nutrients among the study sites. Indeed, the content of P.org, RAL and host age were among the most important drivers of fungal community structure, explaining 3.8%, 3.5% and 3.5% of the total variance, respectively. Similarly, Coince et al. (2013) and Glassman et al. (2017) also observed the strong influence of phosphorus content on fungal community composition. As fungal communities are major actors of the soil phosphorus biogeochemical cycles, the composition of these communities is sensitive to the concentration of phosphorus in their environment (Fabiańska et al., 2019; Hartman and Tringe, 2019). The richness of the aboveground vegetation as well as its age have also been reported as strong drivers of fungal community composition (Canini et al., 2019; Glassman et al., 2017; Tedersoo et al., 2020; Urbanová et al., 2015; Wagner et al., 2016).

Finally, despite being significant, it is surprising that the host vitality did not explain more than 1.2% of the variance in fungal community assembly. Halmschlager and Kowalski (2004) found similar results when looking at the microbial communities associated with the roots of healthy and decaying oak trees, where species composition varied greatly between sites but much less between living and dead roots from the same site. Similarly, looking at the fungal communities present in the stem of five invasive plant species, Raghavendra et al. (2017) did not find any effect between healthy and decaying plants. On the contrary, Steinrucken et al. (2016) found significant differences in fungal communities associated with the roots of healthy and decaying trees, but not between study sites. These contrasting results suggest that the influence of the host vitality in shaping the associated microbial communities might be host species-dependent and need further investigation. Finally, the minor effect of the host vitality observed on fungal community structure in our study might be due to our experimental approach, in which we did not investigate the kinetics of colonisation, from healthy to decaying host beech. Although it is nearly impossible to predict the fate of healthy trees in forests, an experimental approach in a greenhouse would deepen our understanding of how fungal communities influence beech vitality under a controlled water stress.

### Combined influence of water stress and host vitality on fungal diversity, community composition, and assembly processes

We observed a significant increase of the fungal richness between healthy and decaying beech hosts, without any effect of the age of the host, when looking at the diversity of the fungal communities associated with healthy and decaying beech trees across a water stress gradient. While not observed here, the influence of climate warming on fungal richness and diversity has already been reported, suggesting distinct susceptibility of certain fungal communities to water stress (Hopkins et al., 2018; Preece et al., 2019; Schmidt et al., 2018). The decay of beech trees could act as a disturbance, generating new habitats and thus promoting fungal diversity. Nevertheless, investigations related to the influence of water stress on the richness and diversity of fungal communities remain to be deepened as different studies are showing contrasting trends (Anthony et al., 2021; Lozano et al., 2014; Zhou et al., 2020).

Our results also indicated the presence of a core microbial community that colonises the root system of beech trees across the water stress gradient regardless of their vitality. Among them we find the fungal EcM genera *Cenoccocum* and *Tomentella*, as well as the saprotrophs *Cladophialophora* and *Mortierella*. These fungal genera, and particularly *Cenoccocum* and *Mortierella,* have already been described as ubiquitous and among the most abundant taxa within the soil (Buée et al., 2009; Cairney and Chambers, 1999). In addition, members of the genus *Cenoccocum* have already been detected in water-depleted environments, and our results confirm their ability to survive in dry conditions (Fernandez et al., 2013; Lehto and Zwiazek, 2011). On the other hand, we were able to identify specific fungi associated with either a water stress gradient or with the vitality of the beech host, but none of the species was common for both conditions. Saprotrophic fungi were significantly associated with decaying beech while EcM were more abundant in healthy hosts. This seems reasonable because saprotrophs may profit from using the decaying roots as carbon source, whereas EcM fungi may suffer from reduced carbon supply of decaying host trees. Interestingly, Jaeger et al. (2024) found the opposite pattern in a mesocosm study looking at microbial communities and trophic modes in soils around Scots pine saplings during drought-induced tree mortality. They found an increased abundance of EcM fungi around non-vital and dead pine saplings as compared to vital ones. These discrepancies may be explained by the fact that Jaeger et al. (2024) studied fungal communities in soils and not those associated with tree roots. As already suggested by these authors, an increase in EcMs in soils could result from an increase in spore production to survive unfavourable conditions and to build up a spore bank that will allow rapid recolonisation of roots under more benign conditions. The three saprotrophs *Pezicula radicicola*, *Hymenopellis radicata* and *Camposporium appendiculatum* were significantly associated with decaying beech roots. Halmschlager and Kowalski (2004) also observed the presence of *Pezicula radicicola* associated with decaying oaks and suggested that it may act as contributing factors in the oak decline syndrome.

As the vitality of the host was only a minor factor in the fungal structure and only three fungal saprotrophs, but no beech pathogens were significantly associated with decaying beech hosts, it is difficult to believe that these fungal species were involved in beech decay, which is more likely to be caused by water stress. Yet, *Hymenopellis radicata* has already been described as a parasite of beech roots (Breitenbach & Kränzlin, 2005), and these saprotrophs could decompose roots for carbon gain, contributing to the decline of the beech host. However, additional experiments are required to validate these hypotheses.

Even though we did not observe an influence of water stress on fungal guilds, we identified fungal species significantly associated with beech root systems across a water stress gradient, suggesting a species-specific response to drought. The endophytic fungus *Oidiodendron sp.* and the EcM *Sebacina sp.* and *Lactarius blennius* were detected in the driest sites, while *Mortierella amoeboidea*, *Idriella* sp. and *Ilyonectria mors-panacis* were observed in the wettest ones. To our knowledge, it is the first time that these fungi are reported as being associated with beech root systems under water stress. Noteworthy, *Lactarius blennius* is a beech-specific fungal species (Breitenbach & Kränzlin, 2005), and it would be of great interest to investigate its interactions with its host under water stress in a more controlled environmental set up. Yet, the detection of these fungal communities does not inform us about their activity, and further work should include gene expression analyses to decipher the role of the fungal communities towards drought and host vitality.

As we could not assert the liability of the three fungal saprotrophs in the decay of their host tree, we wondered if the beech vitality combined with water stress exhibited a more general influence on the fungal community assembly process that could not be captured by variance partition. We observed that the assembly process of these fungal communities was significantly altered by the host vitality across a water stress gradient, but in a guild-dependent manner. The assembly process of EcM, saprotrophic and pathogenic communities associated with decaying beech were not influenced by the water stress gradient. These findings suggest that the assembly of fungal communities present in decaying hosts is very stable when facing environmental disturbances. Moreover, this stable assembly supports the findings that fungal communities do not necessarily recover from environmental stress when a certain threshold is crossed (Philippot et al., 2021). On the contrary, in the roots of healthy hosts, we observed an increase in the stochasticity of EcM and saprotrophs across increasing water stress, while determinism increased in pathogens. The patterns of the assembly process of EcM and saprotrophs support the “cry for help” hypothesis when plants are facing stress, changing their root exudation profiles and changing the regulation of their immune defence to modulate the recruitment of microbial partners (Hassani et al., 2018; Mendes et al., 2013; Rolfe et al., 2019). On the contrary, the increase in deterministic assembly for pathogen communities as water stress increases supports the concept that host colonisation is undertaken by more and more specialised pathogens, ending with the decline of the tree.

## CONCLUSION

In summary, we showed that drought related parameters (IDI and Deep soil IDI), and host health are altering the fungal community composition, triggering the assembly of specific fungal taxa likely more tolerant to drought. As fungal communities are involved in biogeochemical cycles and by extension, in host physiology, the alteration of these communities might have important effects on forest resilience under climate change. Therefore, it is crucial to further deepen our understanding of how fungal communities modulate the vitality of their tree hosts and how resilient they are when facing environmental disturbance to help manage our future forests.

## Acknowledgements

We are grateful to Mireia Gomez-Gallego for her assistance in statistical modelling and to Jean-Claude Walser of the Genetic Diversity Centre (GDC) of the ETH Zurich for the processing of fungal community sequences. We would like to thank Felix Gugerli and Christian Rellstab for their contribution to the study design and helpful discussions throughout the project. This research was supported by the Laboratory of Excellence ARBRE (ANR-11-LABX-0002-01) and the SwissForestLab.

## Author Contributions

MP, BD and AK designed and coordinated the research and the experimental design. Sampling was realised by RG, SP and RK, and RG performed tree-ring counting. DNA extractions and ITS amplifications were done by RG and SP. AB and LW provided the topographic and environmental data respectively. Data analyses were performed by FF, MP, BD, and AK. FF, MP, BD, and AK wrote the manuscript. All authors revised the manuscript and approved its final version.

## Availability of data and materials

Raw data were deposited in the NCBI Sequence Read Archive (SRA) with the accession PRJNA1236030.

## Consent for publication

Not applicable

## Ethics approval and consent to participate

Not applicable

## REFERENCES

Acosta-Martínez, V., Cotton, J., Gardner, T., Moore-Kucera, J., Zak, J., Wester, D., Cox, S., 2014. Predominant bacterial and fungal assemblages in agricultural soils during a record drought/heat wave and linkages to enzyme activities of biogeochemical cycling. Applied Soil Ecology 84, 69–82. 10.1016/j.apsoil.2014.06.005

Anthony, M.A., Knorr, M., Moore, J.A.M., Simpson, M., Frey, S.D., 2021. Fungal community and functional responses to soil warming are greater than for soil nitrogen enrichment. Elementa: Science of the Anthropocene 9, 000059. 10.1525/elementa.2021.000059

Bardgett, R.D., Freeman, C., Ostle, N.J., 2008. Microbial contributions to climate change through carbon cycle feedbacks. The ISME Journal 2, 805–814. 10.1038/ismej.2008.58

Beckers, B., Op De Beeck, M., Weyens, N., Boerjan, W., Vangronsveld, J., 2017. Structural variability and niche differentiation in the rhizosphere and endosphere bacterial microbiome of field-grown poplar trees. Microbiome 5, 25. 10.1186/s40168-017-0241-2

Birnbaum, C., Hopkins, A.J.M., Fontaine, J.B., Enright, N.J., 2019. Soil fungal responses to experimental warming and drying in a Mediterranean shrubland. Science of The Total Environment 683, 524–536. 10.1016/j.scitotenv.2019.05.222

Bogeat-Triboulot, M.-B., Bartoli, F., Garbaye, J., Marmeisse, R., Tagu, D., 2004. Fungal ectomycorrhizal community and drought affect root hydraulic properties and soil adherence to roots of *Pinus pinaster* seedlings. Plant Soil 267, 213–223. 10.1007/s11104-005-5349-7

Bonito, G., Reynolds, H., Robeson, M.S., Nelson, J., Hodkinson, B.P., Tuskan, G., Schadt, C.W., Vilgalys, R., 2014. Plant host and soil origin influence fungal and bacterial assemblages in the roots of woody plants. Mol Ecol 23, 3356–3370. 10.1111/mec.12821

Borcard, D., Gillet, F., Legendre, P., 2011. Numerical Ecology with R. Springer, New York, NY. 10.1007/978-1-4419-7976-6

Buée, M., Reich, M., Murat, C., Morin, E., Nilsson, R.H., Uroz, S., Martin, F., 2009. 454 Pyrosequencing analyses of forest soils reveal an unexpectedly high fungal diversity. New Phytologist 184, 449–456. 10.1111/j.1469-8137.2009.03003.x

Cáceres, M.D., Legendre, P., 2009. Associations between species and groups of sites: indices and statistical inference. Ecology 90, 3566–3574. 10.1890/08-1823.1

Cairney, J.W.G., Chambers, S.M. (Eds.), 1999. Ectomycorrhizal Fungi Key Genera in Profile. Springer, Berlin, Heidelberg. 10.1007/978-3-662-06827-4

Canini, F., Zucconi, L., Pacelli, C., Selbmann, L., Onofri, S., Geml, J., 2019. Vegetation, pH and Water Content as Main Factors for Shaping Fungal Richness, Community Composition and Functional Guilds Distribution in Soils of Western Greenland. Front. Microbiol. 10, 2348. 10.3389/fmicb.2019.02348

Castaño, C., Berlin, A., Brandström Durling, M., Ihrmark, K., Lindahl, B.D., Stenlid, J., Clemmensen, K.E., Olson, Å., 2020. Optimized metabarcoding with Pacific biosciences enables semi-quantitative analysis of fungal communities. New Phytologist 228, 1149– 1158. 10.1111/nph.16731

Castro, H.F., Classen, A.T., Austin, E.E., Norby, R.J., Schadt, C.W., 2010. Soil Microbial Community Responses to Multiple Experimental Climate Change Drivers. Appl Environ Microbiol 76, 999–1007. 10.1128/AEM.02874-09

Clemmensen, K.E., Ihrmark, K., Durling, M.B., Lindahl, B.D., 2016. Sample Preparation for Fungal Community Analysis by High-Throughput Sequencing of Barcode Amplicons, in: Martin, F., Uroz, S. (Eds.), Microbial Environmental Genomics (MEG), Methods in Molecular Biology. Springer, New York, NY, pp. 61–88. 10.1007/978-1-4939-3369-3_4

Coince, A., Caël, O., Bach, C., Lengellé, J., Cruaud, C., Gavory, F., Morin, E., Murat, C., Marçais, B., Buée, M., 2013. Below-ground fine-scale distribution and soil versus fine root detection of fungal and soil oomycete communities in a French beech forest. Fungal Ecology 6, 223–235. 10.1016/j.funeco.2013.01.002

Coince, A., Cordier, T., Lengellé, J., Defossez, E., Vacher, C., Robin, C., Buée, M., Marçais, B., 2014. Leaf and Root-Associated Fungal Assemblages Do Not Follow Similar Elevational Diversity Patterns. PLoS ONE 9, e100668. 10.1371/journal.pone.0100668

Cregger, M.A., Veach, A.M., Yang, Z.K., Crouch, M.J., Vilgalys, R., Tuskan, G.A., Schadt, C.W., 2018. The Populus holobiont: dissecting the effects of plant niches and genotype on the microbiome. Microbiome 6, 31. 10.1186/s40168-018-0413-8

Dai, A., 2013. Increasing drought under global warming in observations and models. Nature Clim Change 3, 52–58. 10.1038/nclimate1633

Delgado-Baquerizo, M., Maestre, F.T., Reich, P.B., Trivedi, P., Osanai, Y., Liu, Y., Hamonts, K., Jeffries, T.C., Singh, B.K., 2016. Carbon content and climate variability drive global soil bacterial diversity patterns. Ecological Monographs 86, 373–390. 10.1002/ecm.1216

Dormann, C.F., Elith, J., Bacher, S., Buchmann, C., Carl, G., Carré, G., Marquéz, J.R.G., Gruber, B., Lafourcade, B., Leitão, P.J., Münkemüller, T., McClean, C., Osborne, P.E., Reineking, B., Schröder, B., Skidmore, A.K., Zurell, D., Lautenbach, S., 2013. Collinearity: a review of methods to deal with it and a simulation study evaluating their performance. Ecography 36, 27–46. 10.1111/j.1600-0587.2012.07348.x

Dufrêne, M., Legendre, P., 1997. Species assemblages and indicator species: the need for a flexible asymmetrical approach. Ecological Monographs 67, 345–366. 10.1890/0012-9615(1997)067[0345:SAAIST]2.0.CO;2

Fabiańska, I., Sosa-Lopez, E., Bucher, M., 2019. The role of nutrient balance in shaping plant root-fungal interactions: facts and speculation. Current Opinion in Microbiology 49, 90–96. 10.1016/j.mib.2019.10.004

Fernandez, C.W., McCormack, M.L., Hill, J.M., Pritchard, S.G., Koide, R.T., 2013. On the persistence of Cenococcum geophilum ectomycorrhizas and its implications for forest carbon and nutrient cycles. Soil Biology and Biochemistry 65, 141–143. 10.1016/j.soilbio.2013.05.022

Fu, W., Chen, B., Rillig, M.C., Jansa, J., Ma, W., Xu, C., Luo, W., Wu, Honghui, Hao, Z., Wu, Hui, Zhao, A., Yu, Q., Han, X., 2022. Community response of arbuscular mycorrhizal fungi to extreme drought in a cold-temperate grassland. New Phytologist 234, 2003–2017. 10.1111/nph.17692

Gárate-Escamilla, H., Hampe, A., Vizcaíno-Palomar, N., Robson, T.M., Benito Garzón, M., 2019. Range-wide variation in local adaptation and phenotypic plasticity of fitness-related traits in *Fagus sylvatica* and their implications under climate change. Global Ecol Biogeogr 28, 1336–1350. 10.1111/geb.12936

Gargallo-Garriga, A., Preece, C., Sardans, J., Oravec, M., Urban, O., Peñuelas, J., 2018. Root exudate metabolomes change under drought and show limited capacity for recovery. Sci Rep 8, 12696. 10.1038/s41598-018-30150-0

Gehring, C.A., Sthultz, C.M., Flores-Rentería, L., Whipple, A.V., Whitham, T.G., 2017. Tree genetics defines fungal partner communities that may confer drought tolerance. Proc. Natl. Acad. Sci. U.S.A. 114, 11169–11174. 10.1073/pnas.1704022114

Glassman, S.I., Wang, I.J., Bruns, T.D., 2017. Environmental filtering by PH and soil nutrients drives community assembly in fungi at fine spatial scales. Molecular Ecology 26, 6960– 6973. 10.1111/mec.14414

Hagedorn, F., Joseph, J., Peter, M., Luster, J., Pritsch, K., Geppert, U., Kerner, R., Molinier, V., Egli, S., Schaub, M., Liu, J.-F., Li, M., Sever, K., Weiler, M., Siegwolf, R.T.W., Gessler, A., Arend, M., 2016. Recovery of trees from drought depends on belowground sink control. Nature Plants 2, 16111. 10.1038/nplants.2016.111

Halmschlager, E., Kowalski, T., 2004. The mycobiota in nonmycorrhizal roots of healthy and declining oaks. Can. J. Bot. 82, 1446–1458. 10.1139/b04-101

Hari, Vittal, Oldrich Rakovec, Yannis Markonis, Martin Hanel, and Rohini Kumar. 2020. ‘Increased Future Occurrences of the Exceptional 2018–2019 Central European Drought under Global Warming’. Scientific Reports 10 (1): 12207. 10.1038/s41598-020-68872-9.

Hartman, K., Tringe, S.G., 2019. Interactions between plants and soil shaping the root microbiome under abiotic stress. Biochemical Journal 476, 2705–2724. 10.1042/BCJ20180615

Hartmann, M., Brunner, I., Hagedorn, F., Bardgett, R.D., Stierli, B., Herzog, C., Chen, X., Zingg, A., Graf-Pannatier, E., Rigling, A., Frey, B., 2017. A decade of irrigation transforms the soil microbiome of a semi-arid pine forest. Mol Ecol 26, 1190–1206. 10.1111/mec.13995

Hartmann, Henrik, Catarina F. Moura, William R. L. Anderegg, Nadine K. Ruehr, Yann Salmon, Craig D. Allen, Stefan K. Arndt, et al. 2018. ‘Research Frontiers for Improving Our Understanding of Drought-induced Tree and Forest Mortality’. New Phytologist 218 (1): 15–28. 10.1111/nph.15048.

Hassani, M.A., Durán, P., Hacquard, S., 2018. Microbial interactions within the plant holobiont. Microbiome 6, 58. 10.1186/s40168-018-0445-0

Hendershot, J.N., Read, Q.D., Henning, J.A., Sanders, N.J., Classen, A.T., 2017. Consistently inconsistent drivers of microbial diversity and abundance at macroecological scales. Ecology 98, 1757–1763. 10.1002/ecy.1829

Hoffman, G.E., Schadt, E.E., 2016. variancePartition: interpreting drivers of variation in complex gene expression studies. BMC Bioinformatics 17, 483. 10.1186/s12859-016-1323-z

Hopkins, A.J.M., Ruthrof, K.X., Fontaine, J.B., Matusick, G., Dundas, S.J., Hardy, G.Es., 2018. Forest die-off following global-change-type drought alters rhizosphere fungal communities. Environ. Res. Lett. 13, 095006. 10.1088/1748-9326/aadc19

Jaeger, A.C.H., Hartmann, M., Conz, R.F., Six, J., Solly, E.F., 2024. Drought-induced tree mortality in Scots pine mesocosms promotes changes in soil microbial communities and trophic groups. Applied Soil Ecology 194, 105198. 10.1016/j.apsoil.2023.105198

Lagueux, D., Jumpponen, A., Porras-Alfaro, A., Herrera, J., Chung, Y.A., Baur, L.E., Smith, M.D., Knapp, A.K., Collins, S.L., Rudgers, J.A., 2021. Experimental drought re-ordered assemblages of root-associated fungi across North American grasslands. Journal of Ecology 109, 776–792. 10.1111/1365-2745.13505

Lareen, A., Burton, F., Schäfer, P., 2016. Plant root-microbe communication in shaping root microbiomes. Plant Mol Biol 90, 575–587. 10.1007/s11103-015-0417-8

Lehto, T., Zwiazek, J.J., 2011. Ectomycorrhizas and water relations of trees: a review. Mycorrhiza 21, 71–90. 10.1007/s00572-010-0348-9

Li, P., Xu, T., Hu, Q., Gu, S., Yang, Y., Wang, Z., Deng, X., Wang, B., Li, W., Zhu, Y., 2023. Spatial distribution pattern across multiple microbial groups along an environmental stress gradient in tobacco soil. Ann Microbiol 73, 20. 10.1186/s13213-023-01717-8

Lindahl, B.D., Nilsson, R.H., Tedersoo, L., Abarenkov, K., Carlsen, T., Kjøller, R., Kõljalg, U., Pennanen, T., Rosendahl, S., Stenlid, J., Kauserud, H., 2013. Fungal community analysis by high-throughput sequencing of amplified markers – a user’s guide. New Phytologist 199, 288–299. 10.1111/nph.12243

Lozano, Y.M., Hortal, S., Armas, C., Pugnaire, F.I., 2014. Interactions among soil, plants, and microorganisms drive secondary succession in a dry environment. Soil Biology and Biochemistry 78, 298–306. 10.1016/j.soilbio.2014.08.007

Mangeot-Peter, L., Tschaplinski, T.J., Engle, N.L., Veneault-Fourrey, C., Martin, F., Deveau, A., 2020. Impacts of Soil Microbiome Variations on Root Colonization by Fungi and Bacteria and on the Metabolome of *Populus tremula* × *alba*. Phytobiomes Journal 4, 142–155. 10.1094/PBIOMES-08-19-0042-R

Martin, F., Kohler, A., Murat, C., Veneault-Fourrey, C., Hibbett, D.S., 2016. Unearthing the roots of ectomycorrhizal symbioses. Nat Rev Microbiol 14, 760–773. 10.1038/nrmicro.2016.149

McDowell, N.G., Allen, C.D., Anderson-Teixeira, K., Aukema, B.H., Bond-Lamberty, B., Chini, L., Clark, J.S., Dietze, M., Grossiord, C., Hanbury-Brown, A., Hurtt, G.C., Jackson, R.B., Johnson, D.J., Kueppers, L., Lichstein, J.W., Ogle, K., Poulter, B., Pugh, T.A.M., Seidl, R., Turner, M.G., Uriarte, M., Walker, A.P., Xu, C., 2020. Pervasive shifts in forest dynamics in a changing world. Science 368, eaaz9463. 10.1126/science.aaz9463

Meisner, A., Jacquiod, S., Snoek, B.L., ten Hooven, F.C., van der Putten, W.H., 2018. Drought Legacy Effects on the Composition of Soil Fungal and Prokaryote Communities. Front. Microbiol. 9, 294. 10.3389/fmicb.2018.00294

Mendes, R., Garbeva, P., Raaijmakers, J.M., 2013. The rhizosphere microbiome: significance of plant beneficial, plant pathogenic, and human pathogenic microorganisms. FEMS Microbiol Rev 37, 634–663. 10.1111/1574-6976.12028

Moore, J.A.M., Anthony, M.A., Pec, G.J., Trocha, L.K., Trzebny, A., Geyer, K.M., Van Diepen, L.T.A., Frey, S.D., 2021. Fungal community structure and function shifts with atmospheric nitrogen deposition. Global Change Biology 27, 1349–1364. 10.1111/gcb.15444

Morella, N.M., Weng, F.C.-H., Joubert, P.M., Metcalf, C.J.E., Lindow, S., Koskella, B., 2020. Successive passaging of a plant-associated microbiome reveals robust habitat and host genotype-dependent selection. Proc. Natl. Acad. Sci. U.S.A. 117, 1148–1159. 10.1073/pnas.1908600116

Nguyen, L.T.T., Osanai, Y., Anderson, I.C., Bange, M.P., Tissue, D.T., Singh, B.K., 2018. Flooding and prolonged drought have differential legacy impacts on soil nitrogen cycling, microbial communities and plant productivity. Plant Soil 431, 371–387. 10.1007/s11104-018-3774-7

Nguyen, N.H., Song, Z., Bates, S.T., Branco, S., Tedersoo, L., Menke, J., Schilling, J.S., Kennedy, P.G., 2016. FUNGuild: An open annotation tool for parsing fungal community datasets by ecological guild. Fungal Ecology 20, 241–248. 10.1016/j.funeco.2015.06.006

Nilsson, R.H., Larsson, K.-H., Taylor, A.F.S., Bengtsson-Palme, J., Jeppesen, T.S., Schigel, D., Kennedy, P., Picard, K., Glöckner, F.O., Tedersoo, L., Saar, I., Kõljalg, U., Abarenkov, K., 2019. The UNITE database for molecular identification of fungi: handling dark taxa and parallel taxonomic classifications. Nucleic Acids Research 47, D259–D264. 10.1093/nar/gky1022

Ning, D., Deng, Y., Tiedje, J.M., Zhou, J., 2019. A general framework for quantitatively assessing ecological stochasticity. Proc. Natl. Acad. Sci. U.S.A. 116, 16892–16898. 10.1073/pnas.1904623116

Patejuk, Katarzyna, Anna Baturo-Cieśniewska, Wojciech Pusz, and Agata Kaczmarek-Pieńczewska. 2022. ‘Biscogniauxia Charcoal Canker—A New Potential Threat for Mid-European Forests as an Effect of Climate Change’. Forests 13 (1):89. 10.3390/f13010089.

Pflug, E.E., Buchmann, N., Siegwolf, R.T.W., Schaub, M., Rigling, A., Arend, M., 2018. Resilient Leaf Physiological Response of European Beech (Fagus sylvatica L.) to Summer Drought and Drought Release. Front. Plant Sci. 9, 187. 10.3389/fpls.2018.00187

Philippot, L., Griffiths, B.S., Langenheder, S., 2021. Microbial Community Resilience across Ecosystems and Multiple Disturbances. Microbiol Mol Biol Rev 85, e00026–20. 10.1128/MMBR.00026-20

Pokhrel, Yadu, Farshid Felfelani, Yusuke Satoh, Julien Boulange, Peter Burek, Anne Gädeke, Dieter Gerten, et al. 2021. ‘Global Terrestrial Water Storage and Drought Severity under Climate Change’. Nature Climate Change 11 (3): 226–33. 10.1038/s41558-020-00972-w.

Põlme, S., Abarenkov, K., Henrik Nilsson, R., Lindahl, B.D., Clemmensen, K.E., Kauserud, H., Nguyen, N., Kjøller, R., Bates, S.T., Baldrian, P., Frøslev, T.G., Adojaan, K., Vizzini, A., Suija, A., Pfister, D., Baral, H.-O., Järv, H., Madrid, H., Nordén, J., Liu, J.-K., Pawlowska, J., Põldmaa, K., Pärtel, K., Runnel, K., Hansen, K., Larsson, K.-H., Hyde, K.D., Sandoval-Denis, M., Smith, M.E., Toome-Heller, M., Wijayawardene, N.N., Menolli, N., Reynolds, N.K., Drenkhan, R., Maharachchikumbura, S.S.N., Gibertoni, T.B., Læssøe, T., Davis, W., Tokarev, Y., Corrales, A., Soares, A.M., Agan, A., Machado, A.R., Argüelles-Moyao, A., Detheridge, A., De Meiras-Ottoni, A., Verbeken, A., Dutta, A.K., Cui, B.-K., Pradeep, C.K., Marín, C., Stanton, D., Gohar, D., Wanasinghe, D.N., Otsing, E., Aslani, F., Griffith, G.W., Lumbsch, T.H., Grossart, H.-P., Masigol, H., Timling, I., Hiiesalu, I., Oja, J., Kupagme, J.Y., Geml, J., Alvarez-Manjarrez, J., Ilves, K., Loit, K., Adamson, K., Nara, K., Küngas, K., Rojas-Jimenez, K., Bitenieks, K., Irinyi, L., Nagy, L.G., Soonvald, L., Zhou, L.-W., Wagner, L., Aime, M.C., Öpik, M., Mujica, M.I., Metsoja, M., Ryberg, M., Vasar, M., Murata, M., Nelsen, M.P., Cleary, M., Samarakoon, M.C., Doilom, M., Bahram, M., Hagh-Doust, N., Dulya, O., Johnston, P., Kohout, P., Chen, Q., Tian, Q., Nandi, R., Amiri, R., Perera, R.H., Dos Santos Chikowski, R., Mendes-Alvarenga, R.L., Garibay-Orijel, R., Gielen, R., Phookamsak, R., Jayawardena, R.S., Rahimlou, S., Karunarathna, S.C., Tibpromma, S., Brown, S.P., Sepp, S.-K., Mundra, S., Luo, Z.-H., Bose, T., Vahter, T., Netherway, T., Yang, T., May, T., Varga, T., Li, W., Coimbra, V.R.M., De Oliveira, V.R.T., De Lima, V.X., Mikryukov, V.S., Lu, Y., Matsuda, Y., Miyamoto, Y., Kõljalg, U., Tedersoo, L., 2020. FungalTraits: a user-friendly traits database of fungi and fungus-like stramenopiles. Fungal Diversity 105, 1–16. 10.1007/s13225-020-00466-2

Preece, C., Farré-Armengol, G., Peñuelas, J., 2020. Drought is a stronger driver of soil respiration and microbial communities than nitrogen or phosphorus addition in two Mediterranean tree species. Science of The Total Environment 735, 139554. 10.1016/j.scitotenv.2020.139554

Preece, C., Farré-Armengol, G., Verbruggen, E., Peñuelas, J., 2021. Interactive effects of soil water content and nutrients on root exudation in two Mediterranean tree species. Soil Biology and Biochemistry 163, 108453. 10.1016/j.soilbio.2021.108453

Preece, C., Verbruggen, E., Liu, L., Weedon, J.T., Peñuelas, J., 2019. Effects of past and current drought on the composition and diversity of soil microbial communities. Soil Biology and Biochemistry 131, 28–39. 10.1016/j.soilbio.2018.12.022

Raghavendra, A.K.H., Bissett, A.B., Thrall, P.H., Morin, L., Steinrucken, T.V., Galea, V.J., Goulter, K.C., Van Klinken, R.D., 2017. Characterisation of above-ground endophytic and soil fungal communities associated with dieback-affected and healthy plants in five exotic invasive species. Fungal Ecology 26, 114–124. 10.1016/j.funeco.2017.01.003

Reinhold-Hurek, B., Bünger, W., Burbano, C.S., Sabale, M., Hurek, T., 2015. Roots Shaping Their Microbiome: Global Hotspots for Microbial Activity. Annu. Rev. Phytopathol. 53, 403–424. 10.1146/annurev-phyto-082712-102342

Rillig, M.C., Ryo, M., Lehmann, A., Aguilar-Trigueros, C.A., Buchert, S., Wulf, A., Iwasaki, A., Roy, J., Yang, G., 2019. The role of multiple global change factors in driving soil functions and microbial biodiversity. Science 366, 886–890. 10.1126/science.aay2832

Rolfe, S.A., Griffiths, J., Ton, J., 2019. Crying out for help with root exudates: adaptive mechanisms by which stressed plants assemble health-promoting soil microbiomes. Current Opinion in Microbiology 49, 73–82. 10.1016/j.mib.2019.10.003

Rousk, J., Bååth, E., Brookes, P.C., Lauber, C.L., Lozupone, C., Caporaso, J.G., Knight, R., Fierer, N., 2010. Soil bacterial and fungal communities across a pH gradient in an arable soil. The ISME Journal 4, 1340–1351. 10.1038/ismej.2010.58

Sasse, J., Martinoia, E., Northen, T., 2018. Feed Your Friends: Do Plant Exudates Shape the Root Microbiome? Trends in Plant Science 23, 25–41. 10.1016/j.tplants.2017.09.003

Schmidt, C.S., Lovecká, P., Mrnka, L., Vychodilová, A., Strejček, M., Fenclová, M., Demnerová, K., 2018. Distinct Communities of Poplar Endophytes on an Unpolluted and a Risk Element-Polluted Site and Their Plant Growth-Promoting Potential In Vitro. Microb Ecol 75, 955–969. 10.1007/s00248-017-1103-y

Schuldt, B., Buras, A., Arend, M., Vitasse, Y., Beierkuhnlein, C., Damm, A., Gharun, M., Grams, T.E.E., Hauck, M., Hajek, P., Hartmann, H., Hiltbrunner, E., Hoch, G., Holloway-Phillips, M., Körner, C., Larysch, E., Lübbe, T., Nelson, D.B., Rammig, A., Rigling, A., Rose, L., Ruehr, N.K., Schumann, K., Weiser, F., Werner, C., Wohlgemuth, T., Zang, C.S., Kahmen, A., 2020. A first assessment of the impact of the extreme 2018 summer drought on Central European forests. Basic and Applied Ecology 45, 86–103. 10.1016/j.baae.2020.04.003

Segata, N., Izard, J., Waldron, L., Gevers, D., Miropolsky, L., Garrett, W.S., Huttenhower, C., 2011. Metagenomic biomarker discovery and explanation. Genome Biol 12, R60. 10.1186/gb-2011-12-6-r60

Steinrucken, T.V., Bissett, A., Powell, J.R., Raghavendra, Anil. K.H., Van Klinken, R.D., 2016. Endophyte community composition is associated with dieback occurrence in an invasive tree. Plant Soil 405, 311–323. 10.1007/s11104-015-2529-y

Taylor, D.L., Walters, W.A., Lennon, N.J., Bochicchio, J., Krohn, A., Caporaso, J.G., Pennanen, T., 2016. Accurate Estimation of Fungal Diversity and Abundance through Improved Lineage-Specific Primers Optimized for Illumina Amplicon Sequencing. Appl Environ Microbiol 82, 7217–7226. 10.1128/AEM.02576-16

Tedersoo, L., Anslan, S., Bahram, M., Drenkhan, R., Pritsch, K., Buegger, F., Padari, A., Hagh-Doust, N., Mikryukov, V., Gohar, D., Amiri, R., Hiiesalu, I., Lutter, R., Rosenvald, R., Rähn, E., Adamson, K., Drenkhan, T., Tullus, H., Jürimaa, K., Sibul, I., Otsing, E., Põlme, S., Metslaid, M., Loit, K., Agan, A., Puusepp, R., Varik, I., Kõljalg, U., Abarenkov, K., 2020. Regional-Scale In-Depth Analysis of Soil Fungal Diversity Reveals Strong pH and Plant Species Effects in Northern Europe. Front. Microbiol. 11, 1953. 10.3389/fmicb.2020.01953

Trivedi, P., Leach, J.E., Tringe, S.G., Sa, T., Singh, B.K., 2020. Plant–microbiome interactions: from community assembly to plant health. Nat Rev Microbiol 18, 607–621. 10.1038/s41579-020-0412-1

Urbanová, M., Šnajdr, J., Baldrian, P., 2015. Composition of fungal and bacterial communities in forest litter and soil is largely determined by dominant trees. Soil Biology and Biochemistry 84, 53–64. 10.1016/j.soilbio.2015.02.011

Uroz, S., Buée, M., Deveau, A., Mieszkin, S., Martin, F., 2016. Ecology of the forest microbiome: Highlights of temperate and boreal ecosystems. Soil Biology and Biochemistry 103, 471–488. 10.1016/j.soilbio.2016.09.006

Veach, A.M., Chen, H., Yang, Z.K., Labbe, A.D., Engle, N.L., Tschaplinski, T.J., Schadt, C.W., Cregger, M.A., 2020. Plant Hosts Modify Belowground Microbial Community Response to Extreme Drought. mSystems 5, e00092–20. 10.1128/mSystems.00092-20

Wagner, M.R., Lundberg, D.S., del Rio, T.G., Tringe, S.G., Dangl, J.L., Mitchell-Olds, T., 2016. Host genotype and age shape the leaf and root microbiomes of a wild perennial plant. Nat Commun 7, 12151. 10.1038/ncomms12151

Waldrop, M.P., Firestone, M.K., 2006. Response of Microbial Community Composition and Function to Soil Climate Change. Microb Ecol 52, 716–724. 10.1007/s00248-006-9103-3

Walthert, L., Ganthaler, A., Mayr, S., Saurer, M., Waldner, P., Walser, M., Zweifel, R., von Arx, G., 2021. From the comfort zone to crown dieback: Sequence of physiological stress thresholds in mature European beech trees across progressive drought. Science of The Total Environment 753, 141792. 10.1016/j.scitotenv.2020.141792

Walthert, L., and Patrick S., 2018. ‘Equations to Compensate for the Temperature Effect on Readings from Dielectric *Decagon MPS-2* and *MPS-6* Water Potential Sensors in Soils’. Journal of Plant Nutrition and Soil Science 181 (5): 749–59. 10.1002/jpln.201700620.

Wickham, H., 2016. ggplot2, Use R! Springer International Publishing, Cham. 10.1007/978-3-319-24277-4

Williams, A., De Vries, F.T., 2020. Plant root exudation under drought: implications for ecosystem functioning. New Phytologist 225, 1899–1905. 10.1111/nph.16223

Winterfeldt, S., Cruz-Paredes, C., Rousk, J., Leizeaga, A., 2024. Microbial resistance and resilience to drought across a European climate gradient. Soil Biology and Biochemistry 199, 109574. 10.1016/j.soilbio.2024.109574

Wullschleger, S.D., Hanson, P.J., 2006. Sensitivity of canopy transpiration to altered precipitation in an upland oak forest: evidence from a long-term field manipulation study. Global Change Biology 12, 97–109. 10.1111/j.1365-2486.2005.001082.x

Yang, Y., Li, T., Wang, Y., Cheng, H., Chang, S.X., Liang, C., An, S., 2021. Negative effects of multiple global change factors on soil microbial diversity. Soil Biology and Biochemistry 156, 108229. 10.1016/j.soilbio.2021.108229

Zhou, Z., Wang, C., Luo, Y., 2020. Meta-analysis of the impacts of global change factors on soil microbial diversity and functionality. Nat Commun 11, 3072. 10.1038/s41467-020-16881-7

Zuo, X., Sun, S., Wang, S., Yue, P., Hu, Y., Zhao, S., Guo, X., Li, X., Chen, M., Ma, X., Qu, H., Hu, W., Zhao, X., Allington, G.R.H., 2023. Contrasting relationships between plant-soil microbial diversity are driven by geographic and experimental precipitation changes. Science of The Total Environment 861, 160654. 10.1016/j.scitotenv.2022.160654

Zuur, A.F., Ieno, E.N., Walker, N., Saveliev, A.A., Smith, G.M., 2009. Mixed effects models and extensions in ecology with R, Statistics for Biology and Health. Springer, New York, NY. 10.1007/978-0-387-87458-6

